# Abdominal symptoms during primary Sjögren’s syndrome: a prospective study

**DOI:** 10.1101/2020.01.08.898379

**Authors:** Simon Parreau, Jérémie Jacques, Stéphanie Dumonteil, Sylvain. Palat, Sophie Geyl, Guillaume Gondran, Holy Bezanahary, Eric Liozon, Julie Azaïs, Stéphanie Colombie, Marie-Odile Jauberteau, Véronique Loustaud-Ratti, Kim-Heang Ly, Anne-Laure Fauchais

## Abstract

**Context:** Abdominal symptoms are poorly documented during primary Sjögren’s syndrome (pSS).

**Objectives:** To describe abdominal symptoms among pSS patients and to assess their association with characteristics of the disease.

**Methods:** One hundred and fifty patients followed at Hospital and University Center of Limoges were prospectively included and were evaluated using a composite global symptom score (GSS) describing abdominal symptoms and their severity. Data concerning the clinical and biological characteristics of the pSS and abdominal disorders were also collected.

**Results:** Ninety-five per cent of pSS patients suffered from abdominal symptoms with a median GSS of 7.5±5.5 points out of 30. More than half of the patients experienced abdominal tension (68%), upper abdominal pain (54%), abdominal discomfort (58%) and/or constipation (54%). Regarding the pSS activity ESSDAI score items, general and central nervous system involvement was associated with a high GSS. Regarding the patients’ symptoms ESSPRI score, there was a positive correlation with the GSS (*p<0.01*). Multivariate analysis showed a statistical association between a high GSS and seronegative status for SSA, gastroparesis and ESSPRI score (*p<0.01* for each one).

**Conclusion:** This study revealed that more than 90% of pSS patients suffered from abdominal symptoms. There is currently no therapeutic recommendation because of the lack of specific study and comprehension of the physiopathological mechanisms involved.

## Introduction

Primary Sjögren’s syndrome (pSS) is a chronic autoimmune disorder characterized by lymphocytic infiltration of the exocrine glands and loss of secretory function with oral and eye dryness. It affects predominantly women with a female/male ratio of 9:1 and the peak frequency of the disease is about age 50 years. The etiology is still not well understood (1).

Abdominal disorders affect approximately 25% of pSS patients but are poorly documented in the literature. Indeed, epidemiological data vary according to the studies because of small number of patients and old classification criteria. Data concerning the prevalence of abdominal symptoms and their associations with clinical and biological pSS-related manifestations are also contradictory (2). This heterogeneity of data led us to conduct a prospective study to describe abdominal symptoms in pSS patients as defined by the new 2016 ACR-EULAR criteria (3) and to assess their relationship with the characteristics of the disease.

## Methods

### Population

From July 2017 to June 2018, we conducted a single-center study at the Hospital and University Center of Limoges. It was a prospective, interventional study, validated by the Committee for the Protection of Individuals and registered on clinical.trials.gov (NCT03157011). All patients (age >18 years) followed in our center for pSS in consultation or hospitalization had a proposal to participate in the study. All of them fulfilled the 2016 ACR-EULAR classification criteria for pSS (3) and tested negative for HIV and hepatitis C virus (HCV). They gave their written consent after receiving information about participating in the study. Patients’ characteristics (age at disease onset, sex, duration of pSS), clinical manifestations of pSS, associated organ-specific autoimmune disorders and treatment were recorded. Activity of pSS measured by ESSDAI score (EULAR Sjögren’s syndrome disease activity index) and pSS’s symptoms measured by ESSPRI score (EULAR Sjogren’s Syndrome Patient Reported Index) were systematically evaluated at the same time as the abdominal score (4).

Hematological, biochemical, and immunological tests included blood count, fibrinogen, C-reactive protein test, renal and hepatic function tests, β2-microglobulin, serum protein electrophoresis, anti-nuclear antibodies (ANA) (indirect immunofluorescence), precipitating antibodies to the extractable nuclear antigens Ro/SSA and La/SSB (enzyme-linked immunosorbent assay), rheumatoid factor (RF) (latex fixation and Waaler–Rose tests), free light chains and complement dosage. Cryoglobulin was evaluated in case of related clinical manifestations and/or diminution of complement. All tests were performed for each patient at the time of diagnosis and during the completion of the abdominal symptoms score.

### Evaluation of abdominal symptoms

All pSS patients were interviewed by a physician using a questionnaire. It concerned ten abdominal symptoms (nausea, vomiting, upper and lower abdominal pain, abdominal discomfort, bloating, diarrhea, constipation, tenesmus, dysuria). Each symptom carried a score from 0 (no symptom) to 3 (severe) and evaluated by professional judgement. A global symptomatic score (GSS), calculated as the addition of all symptom scores, was assigned to each patient (maximum score 30) (5).

Previously diagnosed digestive, pancreatic and hepatic diseases were also systematically recorded by on the medical file as well as the current symptomatic treatment taken for digestive complaints. Previously diagnosis of gastroparesis was based on international clinical guidelines (6) and irritable bowel syndrome on Rome IV classification (7). All the patients did not have a digestive exploration after the questionnaire but only those with alarm signs (endoscopy) or signs evocative of gastroparesis (gastric emptying scintigraphy) according to the French recommendations (8–10).

### Statistical analysis

Quantitative variables are described by their mean ± standard deviation, and qualitative variables by number and percentage. For qualitative variables, a Chi2 or Fisher’s exact test was used to compare the different groups of patients (SSA +/− SSB patients and seronegative patients). For quantitative variables in a normal distribution, the Student’s t-test was used to compare groups of two or more classes. For quantitative variables that do not follow a normal distribution, the Wilcoxon signed-rank test or the Kruskal-Wallis test was used to compare groups of two or more classes. Univariate analyses between different abdominal symptoms and the rest of the variables were performed. Variables with a *p* value less than 0.20 were included in a multivariate logistic model, simplified by a stepwise elimination method (11), so that the final model only includes variables that are significantly associated with the digestive symptoms. Quantitative variables verifying the Logit linearity assumption were incorporated without modification, or otherwise categorized. The relevance of the model was assessed using Pearson’s residual and deviance tests, and its quality with an OCR curve. All statistical analyses were done using R software (version 3.2.2). p Values less than 0.05 were considered significant.

## Results

### Abdominal symptoms

A total of 150 patients (mean age 63±13 y/o, male n=9) with pSS were recruited between July 2017 and June 2018. Ninety-five per cent of patients (n=143) suffered from abdominal symptoms with a median global score of 7.5±5.5 points out of 30. The abdominal complaints collected from the global score are represented according to their degree of severity in Figure 1.

**Figure.**
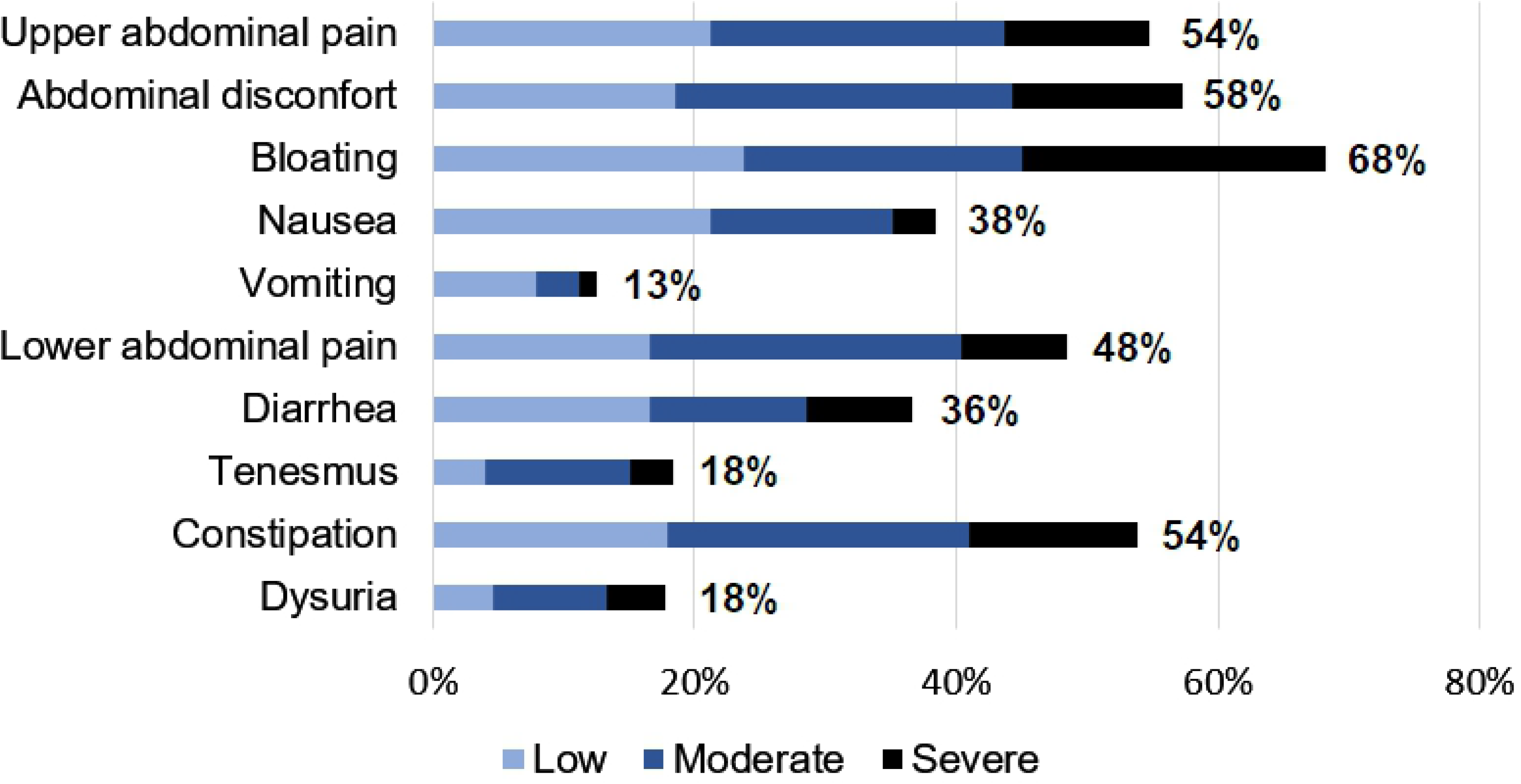

Forty-height percent used at least one treatment for abdominal symptoms in the last 2 weeks. The most highly represented classes were proton pump inhibitors (n=45; 30%), laxatives (n=14; 9%), antispasmodics (n=19; 13%), sodium alginates and/or coals (n=11; 7%), ursodeoxycholic acid (n=6; 4%), prokinetics (n=5; 3%) and anti-diarrheal drugs (n=5; 3%).

### pSS characteristics

Eighty patients (53%) had anti-SSA antibodies and 31 (21%) had anti-SSB antibodies (22%). One hundred and four patients (86%) had a Chisholm score greater than or equal to 3 at the time of diagnosis. None of them had auto-antibodies, symptoms or capillaroscopy evocative of an associated systemic sclerosis. The most common systemic involvements were articular (n=107; 71%), and muscular (n=68; 45%). Twenty-two per cent of patients (n=33) had neurological involvement {central n=20 (13%) and/or peripheral n=24 (16%) with 14 (9%) biopsy-proven small-fiber neuropathies}. Other systemic involvements were hematologic (n=30; 20%), pulmonary (n=17; 11%), cutaneous vasculitis (n=11; 7%) and interstitial nephritis (n=8; 5%) (Supplemental Table 1).

Hydroxychloroquine concerned 46% (n=70) of patients, corticosteroids 23% (n=35), immunosuppressive drugs 14% (n=22), targeted therapies 3% (n=5), intravenous immunoglobulins 5% (n=8). Symptomatic treatment included standard analgesics (n=51; 34%), specific neuropathic pain treatment (n=26; 17%) and nonsteroidal anti-inflammatory drugs for 8% (n=13). Seventeen percent of patients (n=26) used antidepressants and 15% (n=23) benzodiazepines. During the study, the average ESSDAI score was 3.4 ±4.8 points out of 123 and the average ESSPRI score was 17.7±6.2 out of 30.

### Previous digestive disease

Forty-four patients had symptomatic gastroesophageal reflux (29%). Thirteen patients had at least one gastric emptying scintigraphy for clinical signs suggestive of gastroparesis in the cohort (9%) and among them nine had a gastroparesis confirmed. Of these nine patients, three had mild, three moderate and three severe cases of gastroparesis, depending on fixation rate at four hours (6). All patients received medical treatment, three patients were treated with botulinum toxin injections and two underwent endoscopic pylorotomy. Twenty-seven cases of gastritis (18%), mostly atrophic, were reported as well as three cases of associated pernicious anemia (2%). Thirty cases of irritable bowel syndrome were documented (20%). Sixteen patients had enteric endoscopic diverticulosis (11%), which was complicated by diverticulitis in five cases. Three cases of dolichocolons were noted. Sixty-two percent of patients (n=93) had abdominal surgery; appendicectomy (n=44; 29%), cholecystectomy (n=15; 10%) and hysterectomy (n=27; 18%) were the most common.

### Characteristics influencing the abdominal symptoms score

Some characteristics of pSS were associated with a high GSS (Table 1). Regarding the ESSDAI score items, presence of general or central nervous system involvement were associated with a high GSS (*p*=0.0137 and 0.0365 respectively). Regarding the total ESSPRI score, there was a correlation with the GSS of 0.48 (*p*<0.0001). The three items of the ESSPRI score taken separately were also correlated with a high GSS (dryness *p*=0.0014, fatigue *p*<0.0001 and pain *p*<0.0001).

**Table 1.** Association between a high GSS and characteristics of pSS.

|  | High GSS |  | Low GSS |  | *p-value*<sup>a</sup> |
| --- | --- | --- | --- | --- | --- |
|  | n (%) | GSS mean (sd) | n (%) | GSS mean (sd) |  |
| **pSS characteristics** |  |  |  |  |  |
| SSA negative status | 80 (53) | 11.5 (6.1) | 70 (47) | 8.1 (6.0) | <0.0001 |
| Articular involvement | 107 (71) | 10.6 (6.3) | 43 (29) | 7.3 (5.1) | 0.0028 |
| Muscular involvement | 67 (44) | 10.8 (6.4) | 83 (55) | 8.7 (5.9) | 0.0407 |
| **Treatment** |  |  |  |  |  |
| Standard antalgics | 51 (34) | 11.5 (6.6) | 99 (66) | 8.8 (5.9) | 0.0104 |
| Neurotropic antalgics | 26 (17) | 12.0 (6.6) | 124 (83) | 9.2 (6.0) | 0.0397 |

*GSS global symptom score; sd standard deviation; pSS primary Sjögren's syndrome*
<sup>a</sup> For quantitative variables in a normal distribution, the Student's t-test was used to compare groups of two or more classes; for variables that do not follow a normal distribution, the Wilcoxon signed-rank test or the Kruskal-Wallis test was used to compare groups of two or more classes. The value $p < 0.05$ was considered significant

Several digestive antecedents were correlated with a high GSS including hiatal hernia (*p*=0.007), gastroesophageal reflux (*p*=0.001), gastroparesis (*p*=0.0032), atrophic gastritis (*p*=0.0097), irritable bowel syndrome (*p*=0.0002), existence of at least one previous abdominal surgery (*p*=0.0440), and associated pernicious anemia (*p*=0.0306). The consummation of some symptomatic gastrointestinal related treatments was correlated with a high GSS: trimebutine (*p*=0.0047), proton pump inhibitors (*p*=0.0004). In addition, a high GSS was correlated with a higher frequency of endoscopic explorations and gastric emptying scintigraphies after evaluation of the GSS (*p*=0.0037). Concerning pSS specific therapy, none was correlated with a high abdominal symptoms score.

There was no difference between the SSA/SSB positive and seronegative groups with regard to previous digestive diseases and treatment for abdominal symptoms (Supplemental Table 1). On the other hand, serologic status influenced the overall abdominal score as well as prevalence of some abdominal symptoms (Supplemental Table 2). Indeed, the median value for the GSS was significantly higher in seronegative patients (*p*=0.02).

Multivariate analyses of the total GSS (linear regression) and upper gastrointestinal symptoms (logistic regression) *versus* patient characteristics are summarized in Table 2. Multivariate analysis confirmed the statistical association between the GSS total score, seronegative status and ESSPRI score (*p*<0.01, Table 2). There were no items significantly related with lower abdominal symptoms after multivariate analysis. Fatigue and fever of the GSS score were related with the seronegative status (OR=8.45; *p*=0.0152) and a high ESSPRI score (OR=1.31; *p*<0.0001).

**Table 2.** Multivariate analysis investigating effect of GSS on patient characteristics.

| **GSS total** | Coef | 95% CI | *p-value*<sup>a</sup> |
| --- | --- | --- | --- |
| SSA negative status | 2.34 | [0.62-4.06] | 0.00812 |
| Number of past lower abdominal events | 0.74 | [0.23-1.25] | 0.00499 |
| Antecedent of gastroparesis | 4.77 | [1.20-8.35] | 0.00912 |
| ESSPRI score | 0.36 | [0.21-0.51] | <0.0001 |
| **Upper abdominal symptoms** | OR |  |  |
| Seronegative status | 6.01 | [1.52-40.12] | 0.0239 |
| Digestive treatment | 4.64 | [1.37-21.50] | 0.0241 |
| ESSPRI score | 1.11 | [1.02-1.21] | 0.0201 |

*Coef coefficient; CI confidence interval; GSS global symptoms score; OR odds ratio*
<sup>a</sup> $p$ -values for trend are obtained with multivariable linear regression for GSS and logistic regression for upper abdominal symptoms

## Discussion

This study is the first to use the latest criteria of the 2016 ACR-EULAR and to evaluate main abdominal symptoms in order to assess their severity during pSS. Abdominal disorders are rarely reported during pSS and probably underestimated (2). Data in the literature are contradictory in terms of prevalence, with large fluctuations depending on the diagnostic criteria of pSS employed and small number of patients evaluated. Moreover, studies often focus on one type of abdominal involvement (2).

In this prospective study, abdominal signs are frequent and affect most patients with pSS. These symptoms are probably responsible for deterioration in the quality of life of the patients. Indeed Krogh *et al.* have demonstrated that abdominal complaints were a source of impairment of quality of life for pSS patients (12). Despite their frequency, the digestive symptoms are not part of the ESSDAI activity score of pSS. Likewise, the ESSPRI score evaluating the frequent symptoms of pSS does not include the abdominal complaints, which are frequent.

Almost half of the patients use symptomatic digestive treatments. The consumption of symptomatic treatments was correlated with a high abdominal score suggesting that they are not effective and especially unsuitable for digestive involvement of pSS. Otherwise, there was no influence of specific pSS treatments on abdominal symptoms suggesting their ineffectiveness.

Dysfunction of the autonomic nervous system seems to have an important role. Autonomic neuropathy is described during pSS with orthostatic hypotension, urinary retention, segmental anhidrosis and dysfunction of the digestive and urinary systems (13). Muscarinic receptors M3 are located in vascular smooth muscle cells, particularly in gastrointestinal and genitourinary systems as well as exocrine gland cells (14,15). During pSS, it has been found that autoantibodies are directed against muscarinic receptors (14). This suggests that these antibodies may be the cause of autonomic neuropathy and could lead to bladder insufficiency or gastrointestinal disorders (15). A study revealed the involvement of these autoantibodies in the alteration of colonic contraction on an ex-vivo model (16). Intravenous immunoglobulin may be promising for the treatment of immunologically induced digestive or urinary disorders. Smith *et al.* have shown that intravenous immunoglobulin neutralized anti muscarinic M3 receptor antibodies and improved urinary and diarrhea scores for a patient with pSS (17).

Multivariate analysis showed an association between a high abdominal symptom score and antecedent of gastroparesis. Gastroparesis is probably under diagnosed during pSS. We report 69% of confirmed gastroparesis among patients performed an emptying gastric scintigraphy for evocative symptoms. Few studies evaluated gastroparesis during pSS (14,18,19). Kovacs et al. revealed 70% of gastroparesis proven on scintigraphy among 30 pSS patients (14). Another study of 28 pSS patients showed a prevalence of 43% for subjective signs of gastroparesis and 29% for gastroparesis confirmed by octanoate breath tests (18). A retrospective study of 11 patients with pSS-associating confirmed gastroparesis showed that 82% had bloating and abdominal pain (19). The link between gastroparesis and involvement of the autonomic nervous system remains unclear during pSS. The main hypothesis would come from anti-muscarinic receptor type 3 antibodies inhibiting gastric emptying (14). Unfortunately, no study correlated the level of these antibodies and the gastric emptying time.

We acknowledge several limitations. This study was monocentric performed for patients followed in a university hospital center that could induce a selection bias. In addition, there was no control group. We did not systematically investigate for digestive disorder by endoscopy. Regarding the hypothetical link between gastrointestinal involvement and autonomic neuropathy, we did not perform physiological testing and quantitative measurement concerning the autonomic system. In the same way, the detection of antimuscarinic receptor antibodies was not made in our study.

In conclusion, abdominal symptoms, which are often ignored, are common during pSS. They may represent a source of chronic deterioration in quality of life. The neurovegetative system seems particularly involved but further prospective studies are needed. Clinicians must remain alerted to abdominal symptoms and strive to best characterize the type of underlying lesions. Moreover, there is currently no therapeutic recommendation regarding pSS abdominal involvement because of the lack of any specific study and comprehension of the physiopathological mechanisms involved.

## Conflict of Interest

The authors disclose no conflicts of interest

---

**Supplemental Table 1.** Characteristics of patients according to serological status.

|  | SSA+ (n=80) | SSA- (n=70)<br>n (%) | TOTAL (n=150) | *p-value*<sup>a</sup> |
| --- | --- | --- | --- | --- |
| Male | 3 (3.7) | 6 (8.7) | 9 (6.0) | n.s. |
| Age at diagnosis, years | 51.2 (13.4) | 55.0 (11.0) | 53.0 (12.5) | n.s. |
| Mean follow-up, years | 10.3 (7.9) | 9.6 (6.9) | 10.0 (7.4) | n.s. |
| Salivary gland enlargement | 21 (26.3) | 19 (27.1) | 40 (26.5) | n.s. |
| Raynaud's phenomenon | 29 (36.3) | 18 (25.7) | 47 (31.1) | n.s. |
| Articular involvement | 56 (70.0) | 51 (72.9) | 107 (70.9) | n.s. |
| Muscular involvement | 32 (40.0) | 35 (50.0) | 67 (44.4) | n.s. |
| Neurologic involvement | 15 (18.8) | 19 (27.1) | 34 (22.5) | n.s. |
| Cutaneous vasculitis | 9 (11.3) | 1 (1.4) | 10 (6.6) | 0.026 |
| Pulmonary involvement | 11 (13.8) | 5 (7.1) | 16 (10.6) | n.s. |
| Interstitial nephritis | 6 (7.5) | 2 (2.9) | 8 (5.3) | n.s. |
| Hematological involvement | 23 (28.8) | 8 (11.4) | 31 (20.5) | 0.0090 |
| Non-Hodgkin lymphoma | 2 (2.5) | 0 | 2 (1.3) | n.s. |
| Hypergammaglobulinemia | 35 (43.8) | 2 (2.9) | 37 (24.5) | <0.0001 |
| Positive RF | 12 (15.0) | 2 (2.9) | 14 (9.3) | 0.0115 |
| Cryoglobulinemia | 4 (5.0) | 1 (1.4) | 5 (3.3) | n.s. |
| Symptomatic treatment of pSS | 67 (83.7) | 58 (84.0) | 126 (84.0) | n.s. |
| Specific treatment of pSS | 52 (65.0) | 35 (50.7) | 88 (58.7) | n.s. |
| Digestive treatment | 33 (41.2) | 39 (56.5) | 73 (48.7) | n.s. |
| **mean (sd)** |  |  |  |  |
| ESSDAI score | 3.5 (4.9) | 3.2 (4.8) | 3.4 (4.8) | n.s. |
| ESSPRI score | 16.5 (6.4) | 19.1 (5.7) | 17.7 (6.2) | 0.0082 |

*RF rheumatoid factor; pSS primary Sjögren's syndrome; n.s. not significant; sd standard deviation*
<sup>a</sup> For qualitative variables, a Chi-2 or Fisher's exact test was used to compare different groups of patients (SSA+ patients and seronegative patients). For quantitative variables in a normal distribution, the Student's t-test was used to compare groups of two or more classes; for variables that do not follow a normal distribution, the Wilcoxon signed-rank test or the Kruskal-Wallis test was used to compare groups of two or more classes. The value $p < 0.05$ was considered significant.

**Supplemental Table 2.** Comparison of GSS symptoms and serological status.

|  | SSA+ (n=80) | SSA- (n=70)<br>n (%) or mean (sd) | Total (n=150) | *p-value*<sup>a</sup> |
| --- | --- | --- | --- | --- |
| **Upper abdominal symptoms** | 63 (78.8) | 67 (97.1) | 131 (87.3) | 0.0008 |
| Upper abdominal pain | 31 (38.7) | 52 (75.4) | 84 (56.0) | <0.0001 |
| Abdominal discomfort | 40 (50.0) | 47 (68.1) | 88 (58.7) | 0.0253 |
| Bloating | 50 (62.5) | 54 (78.3) | 105 (70.0) | 0.0367 |
| Nausea | 25 (31.2) | 35 (50.7) | 60 (40.0) | 0.0157 |
| Vomiting | 10 (12.5) | 11 (15.9) | 21 (14.0) | n.s. |
| **Lower abdominal symptoms** | 63 (78.8) | 59 (85.5) | 123 (82.0) | n.s. |
| Lower abdominal pain | 35 (43.7) | 39 (56.5) | 75 (50.0) | n.s. |
| Diarrhea | 21 (26.2) | 35 (50.7) | 56 (37.3) | 0.0021 |
| Tenesmus | 14 (17.5) | 13 (18.8) | 28 (18.7) | n.s. |
| Constipation | 42 (52.5) | 40 (58.0) | 82 (54.7) | n.s. |
| Dysuria | 10 (12.5) | 19 (27.5) | 29 (19.3) | 0.0208 |

*sd standard deviation; GSS global symptoms score; n.s. not significant*
<sup>a</sup> For quantitative variables in a normal distribution, the Student's t-test was used to compare groups of two or more classes; for variables that do not follow a normal distribution, the Wilcoxon signed-rank test or the Kruskal-Wallis test was used to compare groups of two or more classes. The value $p < 0.05$ was considered significant

